# Partial Wwox Loss of Function Increases Severity of Murine Sepsis and Neuroinflammation

**DOI:** 10.1101/2025.01.17.633677

**Authors:** Priscilla De La Cruz, Marta Gomes, Angelia Lockett, Amanda Fisher, Todd Cook, Patricia Smith, Christopher Lloyd, Homer L Twigg, Adrian Oblak, C. Marcelo Aldaz, Roberto F. Machado

## Abstract

**Rationale:** WW domain-containing oxidoreductase (*WWOX*) is a gene associated implicated in both neurologic and inflammatory diseases and is susceptible to environmental stressors. We hypothesize partial loss of Wwox function will result in increased sepsis severity and neuroinflammation.

**Methods:** *Wwox ^WT/P47T^* mice, generated by CRISPR/Cas9, and *Wwox ^WT/WT^* mice were treated with intraperitoneal PBS vs LPS (10mg/kg) and euthanized 12 hours post-injection. Open Field Testing (OFT) and Murine Sepsis Severity Scores (MSS) were utilized to measure sickness behavior and sepsis severity, respectively. Brain tissue was analyzed using immunohistochemistry and PCR to measure neuroinflammation and apoptosis.

**Results:** *Wwox ^WT/P47T^* LPS mice demonstrated a more significant response to sepsis with an increase in sickness behavior, sepsis severity, gliosis, and apoptosis compared to *Wwow ^WT/WT^* LPS littermates.

**Conclusions:** Partial loss of Wwox function increases risk for severe sepsis and neuroinflammation. Given the susceptibility of *WWOX* to environmental stressors, this may be a target for future therapeutic interventions.

## Introduction

Sepsis is a life threatening syndrome that results in end-organ damage and occurs in nearly 1 million patients per year and is responsible for more than half of the deaths identified in hospitalized patients^1,2^. Sepsis-associated encephalopathy (SAE), a form of delirium associated with increased neuroinflammation in setting of sepsis, has been identified in up to 70% of patients^3^. Delirium is associated with an increased hospital length of stay, increased 12-month mortality, and long-term neurocognitive deficits^4–6^. There is a need to identify patients at greatest risk for increased sepsis severity and delirium.

WW domain-containing oxidoreductase (*WWOX*) is a gene that has been implicated in both neurologic and inflammatory diseases. *WWOX* is located at the common chromosomal fragile site FRA16D and has multiple roles including cancer pathogenesis and glucose metabolism^7–9^. In addition to being well-known for its role in neurodevelopmental disorders such as spinocerebellar ataxia type 12 (SCAR 12) and *WWox*-related epileptic encephalopathy (WOREE), it is a risk locus in multiple Alzheimer’s Disease (AD) genome wide association studies (GWAS)^10–12^. Interestingly, Wwox expression is correlated to multiple stressors and stimuli including cancer, metabolic syndrome, and oxidative stress^13–15^. For instance, in response to electronic cigarette and tobacco exposure, Wwox expression is reduced in lung tissue and results in an increase in severity of pneumonia-associated acute respiratory distress syndrome (ARDS) and ventilator-associated ARDS^8,16^. Loss of Wwox expression in endothelial cells is associated with increased permeability likely contributing to these findings^16^.

To better understand the effects of Wwox expression, a Proline to Threonine germline mutation was introduced via CRISP/Cas9 knock-in system to generate a *Wwox^P47T/P47T^* mouse model^17^. This mutation impairs the protein-protein interactive properties of Wwox via the WW domains. The heterozygous *Wwox ^WT/P47T^* mice did not differ significantly from the *Wwox ^WT/WT^* littermates, but the *Wwox^P47T/P47T^* mice demonstrated increased sickness behavior, reduced cumulative survival, increased microglia and astrocyte density and activation compared to *Wwox ^WT/WT^* littermate control mice^17^. Gene Set Enrichment Analysis (GSEA) of brain tissue demonstrated increased enrichment of TNF-α signaling via NFKβ pathways, IL-6 JAK Stat 3 signaling, inflammatory response, and apoptosis in the *Wwox^P47T/P47^*^T^ mice compared to the *Wwox ^WT/P47T^* littermates. Based on these findings and given that Wwox expression may be modulated by diverse stimuli, in this work we hypothesized that heterozygous *Wwox ^WT/P47T^* mice will display an increase in sepsis severity, neuroinflammation, and sickness behavior compared to *Wwox ^WT/WT^* littermates.

## Methods

### Animal Model of Sepsis and Behavioral Testing

All experiments were approved by the Institutional Animal Care and Use Committee at Indiana University. *Wwox^P47T/P47T^* mice were developed using CRISPR/Cas9 knock-in system and generously donated by CMarcelo Aldaz^17^. Mice were bred and genotyped using custom Kompetitive Allele Specific PCR genotyping assay(KASP, LGC Biosearch Technologies, UK)^17^. Mice at a mean age of 2.6 months (n=6 male and n=3 female per group) were randomly assigned to receive intraperitoneal (ip) Phosphate Buffered Saline (PBS) vs 10mg/kg Lipopolysaccharide (LPS-EB Ultrapure) (Invivogen, #tlrl-3pelps). Prior to treatment, mice underwent open field testing (OFT), as previously; this is an established procedure to assess neurologic behavior and function^18^. In brief, mice were placed in 30x30cm open field for 10 minutes with video recordings used to obtain measurements of distance traveled and time spent in center of maze using ANY-maze tracking software. The following day, mice were treated with PBS vs LPS and observed using the murine sepsis severity score (MSS) every 2 hours^19^. This scoring system includes measurements of fur quality, activity, respiration status, and mucus membranes. At four hours post-injection, mice repeated the open field test. At 12 hours post-injection, mice were euthanized with collection of brain and plasma.

### Brain Tissue Immunofluorescence Staining

The brains were grossly dissected along the intracerebral fissure with half snap frozen in liquid nitrogen and stored in -80C for protein and RNA studies. The other half was utilized for immunofluorescence and was fixed with 4% paraformaldehyde overnight followed by 30% sucrose treatment for 24 hours. Brains were stored in optimal tissue cutting medium in -80C until sectioning. Coronal sections were obtained at 30 micrometer thickness and stored in cryoprotectant solution (FD NeuroTechnologies, Inch, Cat#PC102). Sections were washed with 0.3% Phosphate Buffered Saline-Triton X-100 (PBST) and treated with 0.3% Hydrogen Peroxide block for 10 minutes. Sections were then washed and incubated with 10% Donkey Serum (Abcam, #NC1697010) for 1 hour at room temperature. Sections were then incubated with primary antibody overnight: Ionized calcium-binding adaptor molecule 1 (Iba1) 1:500 (Synaptic Systems, #234009), Glial Fibrillary Acidic Protein (GFAP) 1:1000 (Invitrogen, #13-0300), CD31 1:1000 (R&D Systems, #AF3628), Fibrinogen (Dako, #A0080), Neuron-Specific Nuclear Protein (NeuN) 1:1000 (Antibodies.com, #A270544), and Cleaved Caspase 3 (CC3) 1:500 (Cell Signaling, #9664). After overnight incubation, sections were washed with PBST, blocked with 10% Donkey Serum and incubated in secondary antibody for 2 hours. Sections were then washed with PBST and mounted on slides using ProLong Gold Antifade Mountant with DNA Stain DAPI (Invitrogen, #P36931). Slides were imaged using Leica Thunder System. Images were analyzed using Imaris (Oxford Instruments).

### Quantitative Polymerase Chain Reaction (PCR)

RNA was extracted from hemibrains using the RNeasy Plus Mini Kit (Qiagen, # 74134) following manufacturer instruction. RNA quality was assessed using Nanodrop One. RNA was converted to cDNA using the High-Capacity cDNA Reverse Transcription Kit (Applied Biosystems, # 4368814). cDNA, primers, RNase-Free Water, and SYBR Green Mix (Applied Biosystems PowerUp SYBR Green Master Mix (Applied Biosystems, # A25742) were combined to measure cytokine expression. The following primers were used: *IL-6* (Forward ‘TCCTACCCCAACTTCCAATGCTC’, Reverse ‘TTGGATGGTCTTGGTCCTTAGCC’), *IL-1β* (Forward ‘CACCTCTCAAGCAGAGCACAG’, Reverse ‘GGGTTCCATGGTGAAGTCAAC’), *TNF-α* (Forward ‘AAATGGGCTCCCTCTCATCAGTTC’, Reverse ‘TCTGCTTGGTGGTTTGCTACGAC’), and *Gapdh*(Forward ‘CATCACTGCCACCCAGAAGACTG’, Reverse ‘ATGCCAGTGAGCTTCCCGTTCAG’). Fold change was calculated using 2^-ΔΔCT^ method and assessed relative to *Gapdh*.

### Cytometric Bead Assay

Plasma was obtained at time of euthanasia and frozen at -80C until needed. Plasma levels of IL-12p70, TNF-α, Interferon-γ, MCP-1, IL-10, and IL6 were measured following manufacturer instructions (BD Biosciences, # 552364).

### Statistical Analysis

Data was analyzed using unpaired sample t-test to compare *Wwox ^WT/P47T^* LPS to *Wwox ^WT/WT^* LPS OFT, IHC, and CBA experiments. Outliers were identified using ROUT method. This resulted in removal of two LPS *Wwox ^WT/P47T^*, one PBS *Wwox ^WT/P47T^*, three LPS *Wwox ^WT/WT^*, and one PBS *Wwox ^WT/WT^*. The risk for mortality was measured using a simple linear regression model and all other analysis was performed using a one-way ANOVA. Data is presented as a mean and standard error of mean (SEM) unless otherwise noted. Data was analyzed using GraphPad Prism.

## Results

### Partial Loss of Wwox Function Increases Sepsis Severity and Sickness Behavior

Sickness behavior and sepsis severity were assessed using behavioral testing and Murine Sepsis Severity scoring (MSS), respectively^19^. This score has been shown to correlate with development of septic shock and mortality in both polymicrobial and LPS models of sepsis^19,20^. Mice completed pre-injection and 6 hours post-injection open field testing to measure distance traveled and time spent in the center of the maze. There was no difference in OFT measurements between the PBS groups. Both genotypes demonstrate sickness behavior with a reduction in distance traveled post-LPS injection (p <0.0001) (**Figure 1A**). The *Wwox ^WT/P47T^* mice demonstrate an LPS response with a reduced time in center post-injection (p=0.0312), and this was not observed in the *Wwox ^WT/WT^* mice (**Figure 1B**). Post-LPS injection mice were evaluated using the Murine Sepsis Severity score. *Wwox ^WT/P47T^* LPS mice demonstrated a rapid onset of severe sepsis compared to *Wwox ^WT/WT^* LPS mice (p= 0.001617) (**Figure 2**).

**Figure 1.**
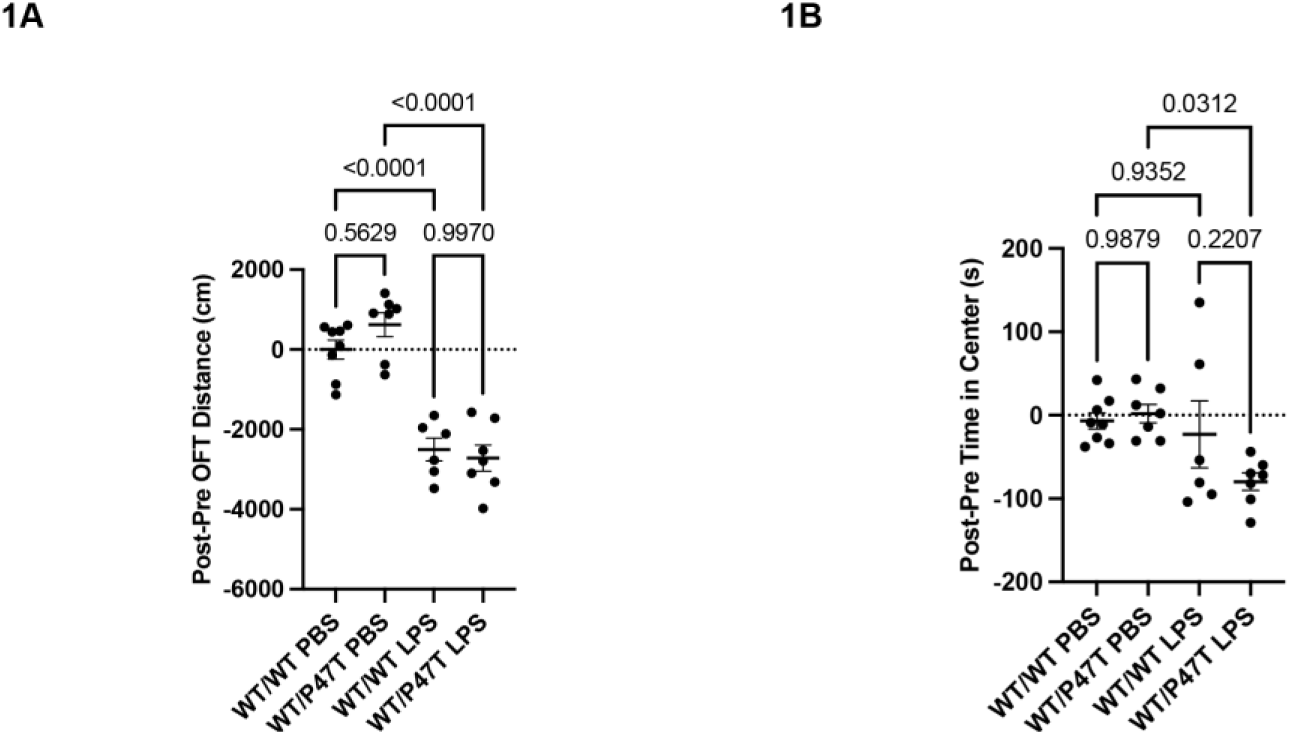
Open Field Test (OFT) demonstrates LPS-induced sickness behavior. Both Wwox genotypes demonstrate an LPS effect with reduced distance traveled post-LPS treatment from pre-treatment baseline. There is no difference in distance between LPS groups (**1A**). Time spent in center is used to measure sickness behavior. There is a difference between the *Wwox ^WT/P47T^* LPS and PBS control groups (p=0.0312) and no difference between the Wwox*™" LPS and PBS control groups (**1B**)

**Figure 2.**
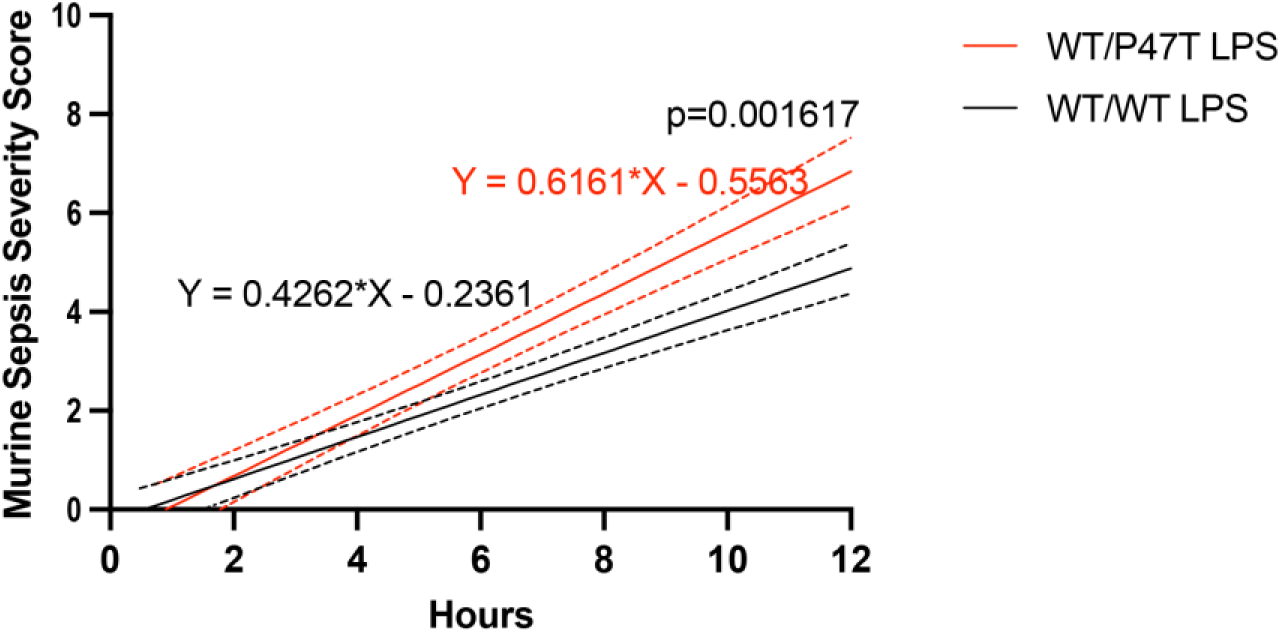
A simple linear regression model comparing *Wwox ^WT/WT^* LPS mice to *Wwox ^WT/P47T^* LPS mice demonstrates an increase in sepsis severity scores at ear1ier times points post-injection in the *Wwox ^WT/P47T^* LPS mice demonstrated by difference in slope that is significantly different (p=0.001617). Solid lines represent mean while dashed lines represent error. *Wwox ^WT/P47T^* mice are represented in red.

### Partial Loss of Wwox Function Increases Neuroinflammation in Experimental Sepsis

Iba1 staining showed no statistically significant difference of frontal cortical microglia density in both PBS groups, while *Wwox ^WT/P47T^* LPS mice exhibited significantly more microglia compared to *Wwox ^WT/WT^* LPS mice (p=0.0282) (**Figure 3A-3B**). This finding was also observed in the hippocampus (p=0.0490) (**Figure 4A-4B**). Next, astrocyte density was measured using GFAP. In the cortex, there was a robust response in the *Wwox ^WT/P47T^* group post-LPS injection with an increase in astrocyte density compared to the PBS *Wwox ^WT/P47T^* group (p=0.0040). There were more cortical astrocytes in the *Wwox ^WT/P47T^* LPS group compared to the *Wwox ^WT/WT^* LPS group (p=0.0005) (**Figure 5A-5B**). There was no difference in astrocyte density in the hippocampus between groups (**Figure 6A-6B**).

**Figure 3.**
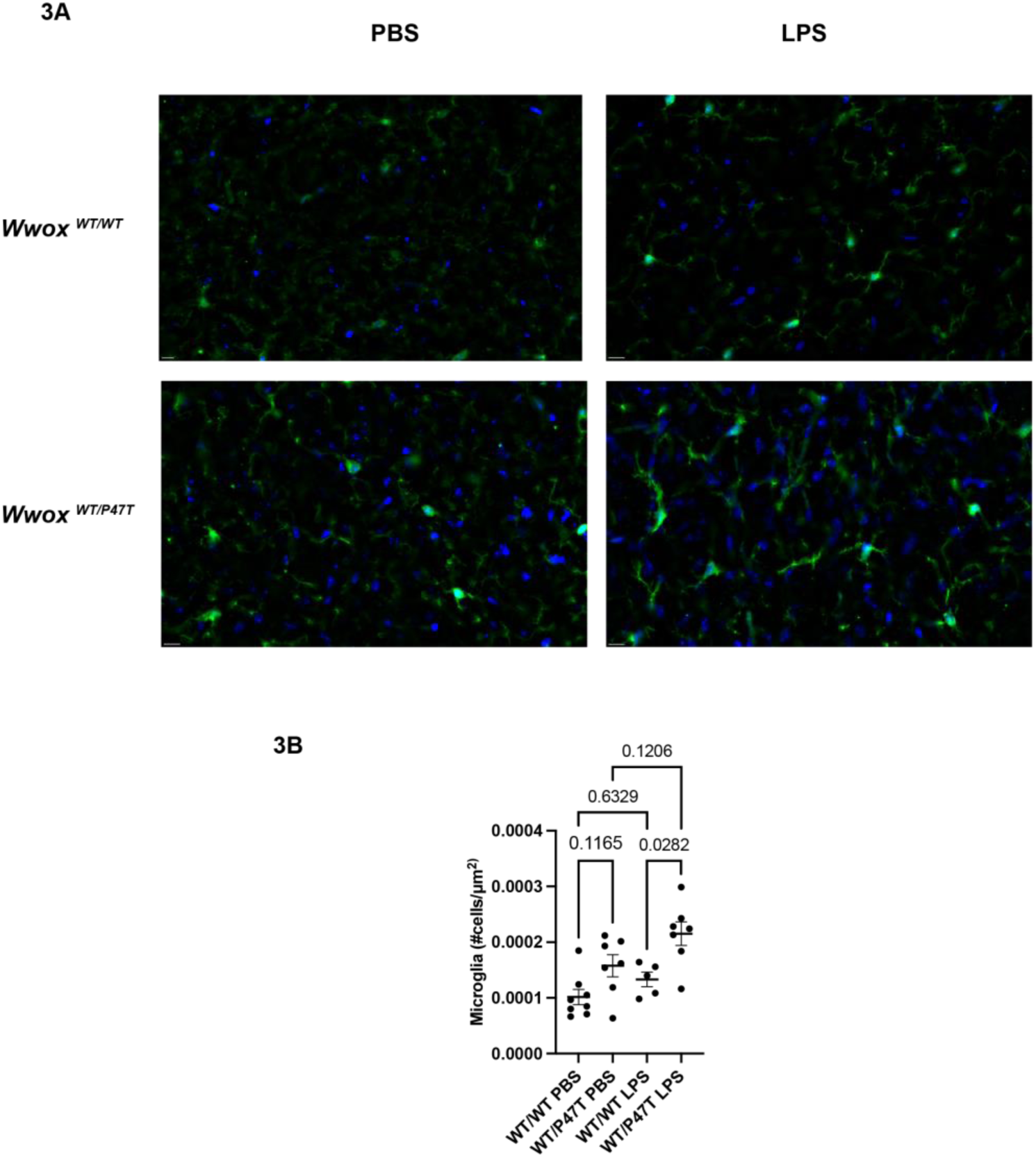
Immunohistochemistry analysis of cortical sections is used to measure microglia (lba1) cell density. Images of microglia are demonstrated with scale bar at 10μm **3A.** one-way ANOVA analysis shows no statistically significant difference in microglia density in thePBS groups and an increased density of microglia in the *Wwox ^WT/P47T^* LPS group compared to the *Wwox ^WT/WT^* LPS group (p=0.0282) **3B.**

**Figure 4.**
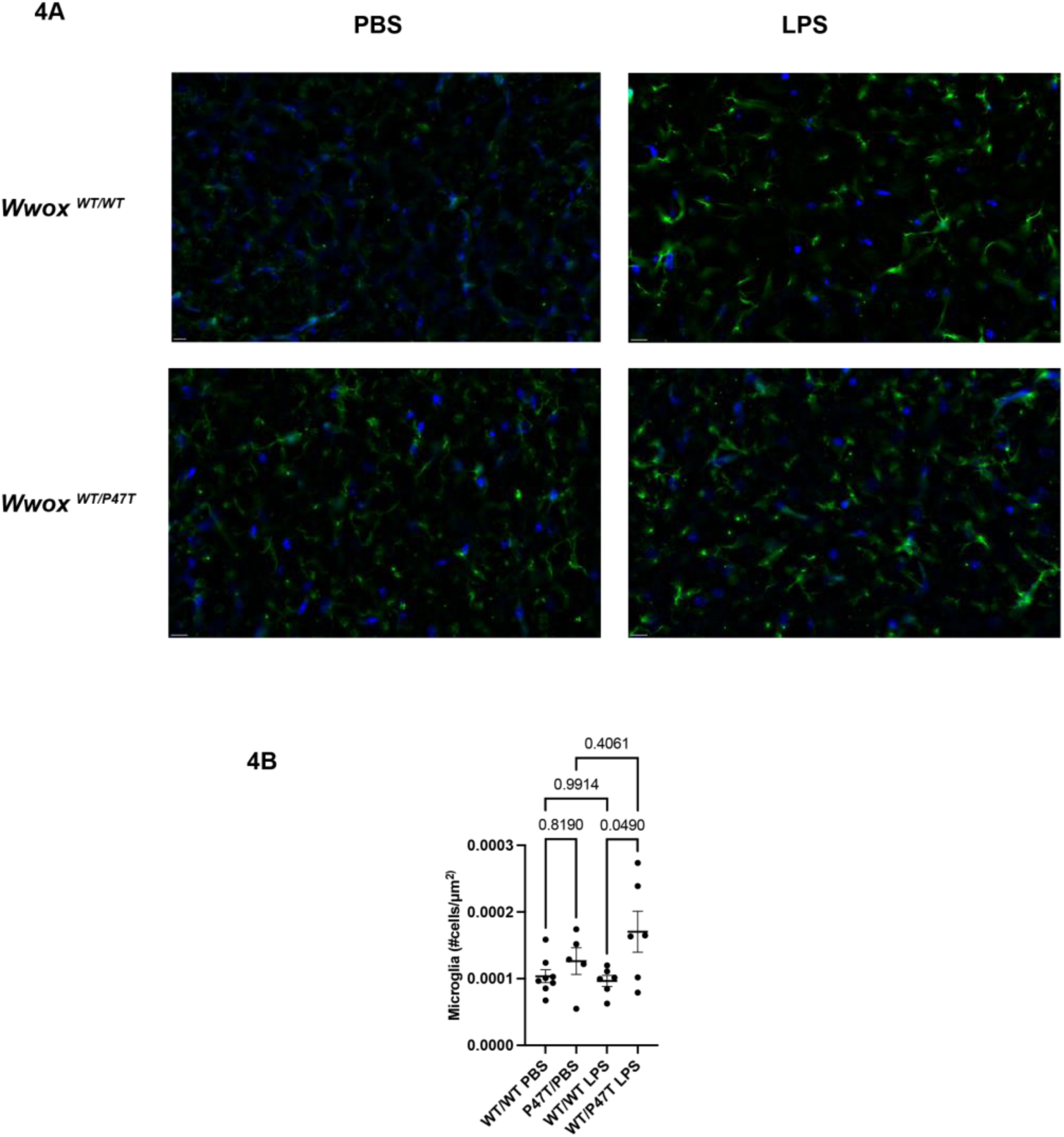
Immunohistochemistry is utilized to measura microglia (lba1) density in the hippocampus. Images of microglia ara demonstrated with scale bar at 10 µm **4A.** One-way ANOVAdemonstrates an increase in microglia density in the *Wwox ^WT/P47T^* LPS mice compared to the *Wwox ^WT/WT^* LPS mice (p=0.0490) **4B.**

**Figure 5.**
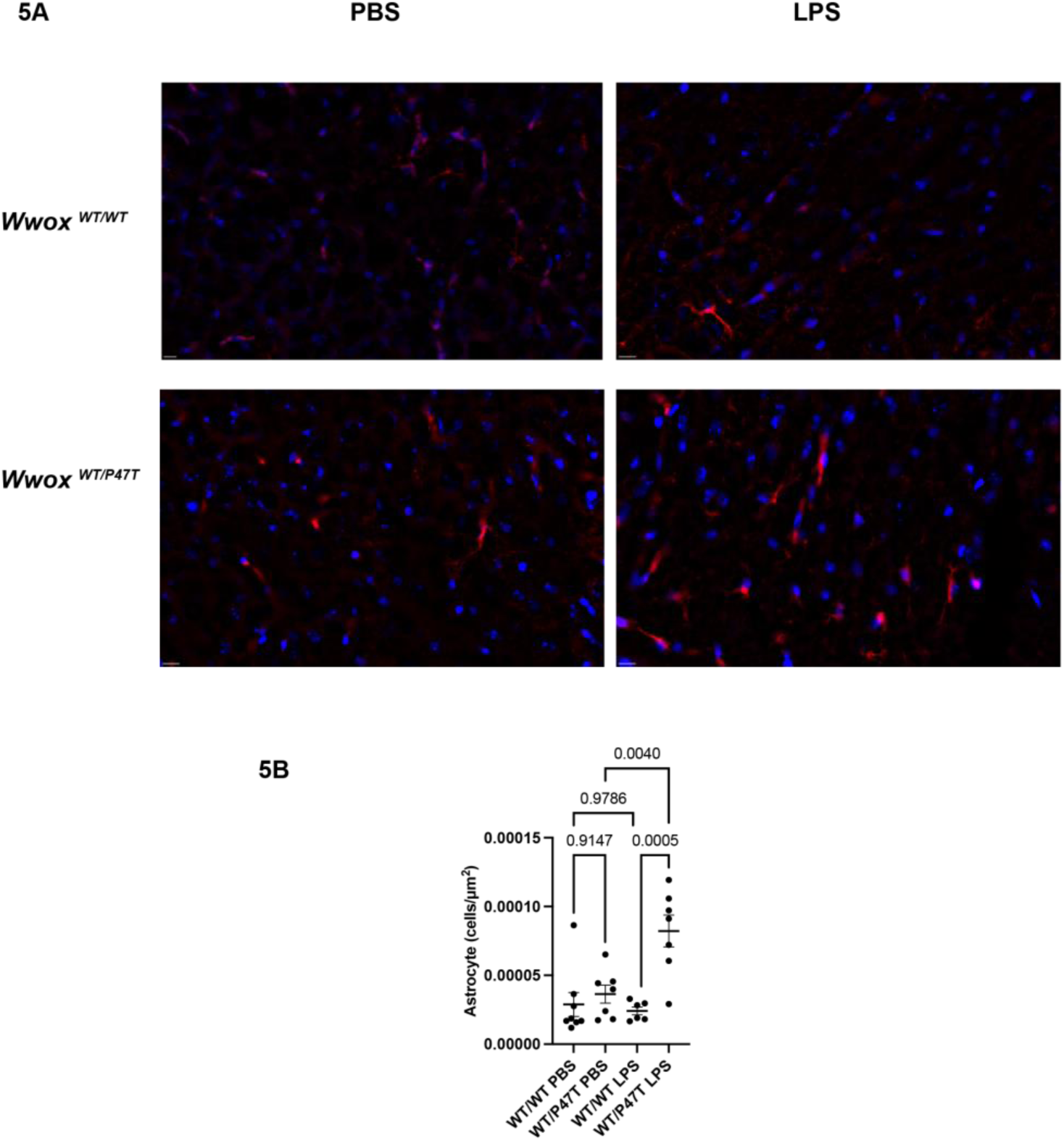
Immunohistochemistry is utilized to measura astrocyte (GFAP) density in cortex. Images ara provided with scale bar at 10μm **5A.** A one-way ANOVA demonstrates an increase in astrocyte density in the *Wwox ^WT/P47T^* LPS mice compared to the *Wwox ^WT/WT^* LPS mice (p=0.0005). There is also an increase in astrocyte density in response to LPS in the *Wwox ^WT/P47T^* group compared to PBS control (p=0.0040) **5B.**

**Figure 6.**
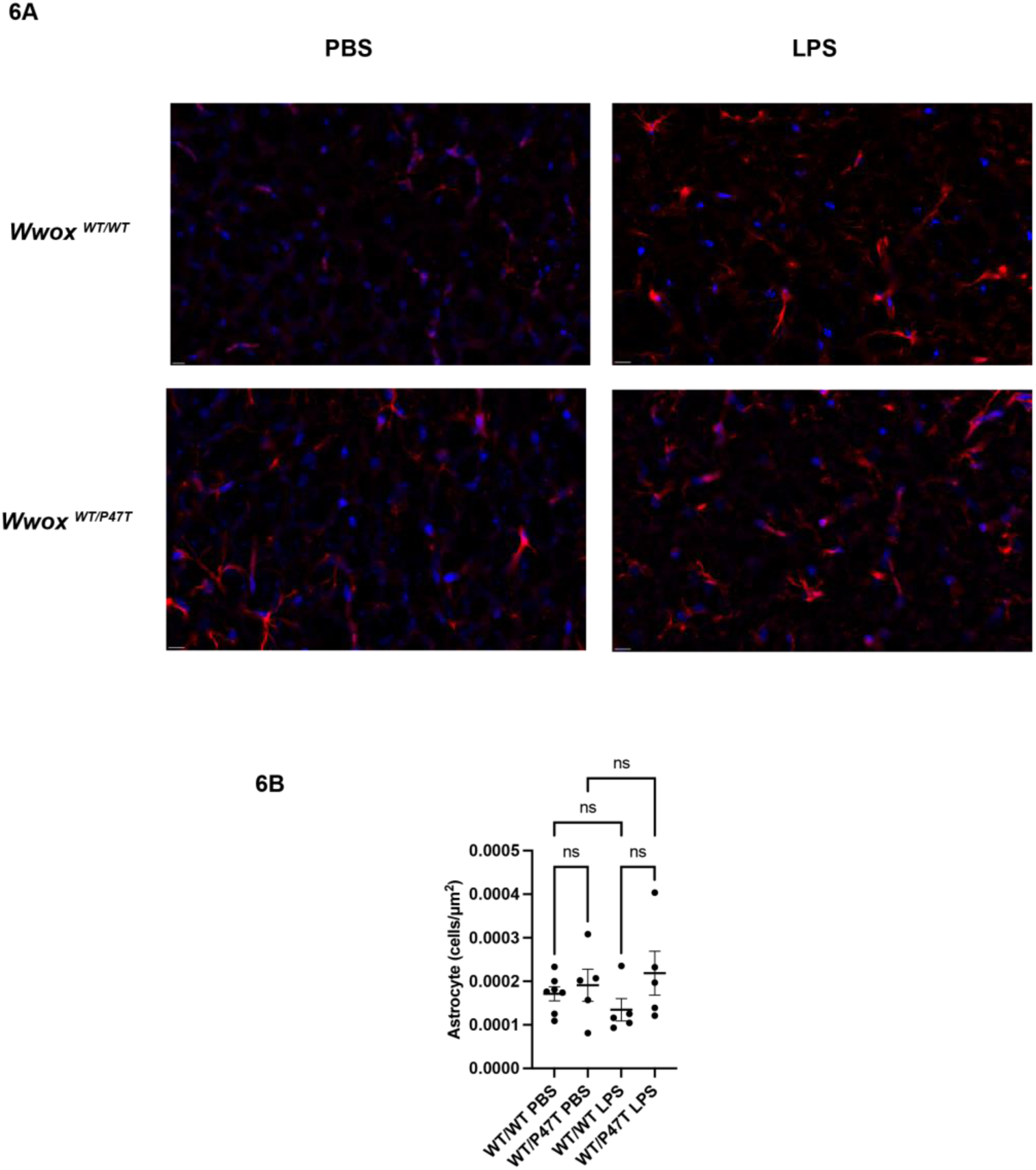
Immunohistochemistry is utilized to measure astrocyte (GFAP) density in the hippocampus. Images are provided with scale barat 10µm **6A.** A one-way ANOVA is used to compare astrocyte density does not demonstrate a significant difference between genotypes post-LPS treatment **6B.**

Quantitative PCR of cortical tissue was used to measure multiple inflammatory markers. Cortex samples from both the *Wwox ^WT/WT^* (p=0.0495) and *Wwox ^WT/P47T^* (p<0.0001) mice displayed evidence of an LPS response in IL-1β levels with an increase in mRNA compared to PBS control groups and a trend towards a more significant response in the *Wwox ^WT/P47T^* LPS group compared to the *Wwox ^WT/WT^* LPS group (p= 0.0837) (**Figure 7A**). The *Wwox ^WT/WT^* group did not show a significant difference in IL-6 mRNA expression with LPS treatment while the *Wwox ^WT/P47T^* mice had an increase in IL-6 with LPS compared to the PBS control (p=0.0303) (**Figure 7B**). Similarly, *the Wwox ^WT/WT^* mice did not demonstrate a difference in TNF-α mRNA levels with LPS. However, the *Wwox ^WT/P47T^* samples showed a significant increase in TNF-α expression observed with LPS treatment compared to PBS control (p=<0.0001) as well as a significant difference in TNF-α levels between the *Wwox ^WT/WT^* LPS and *Wwox ^WT/P47T^* LPS groups (p=0.0016) (**Figure 7C**).

**Figure 7.**
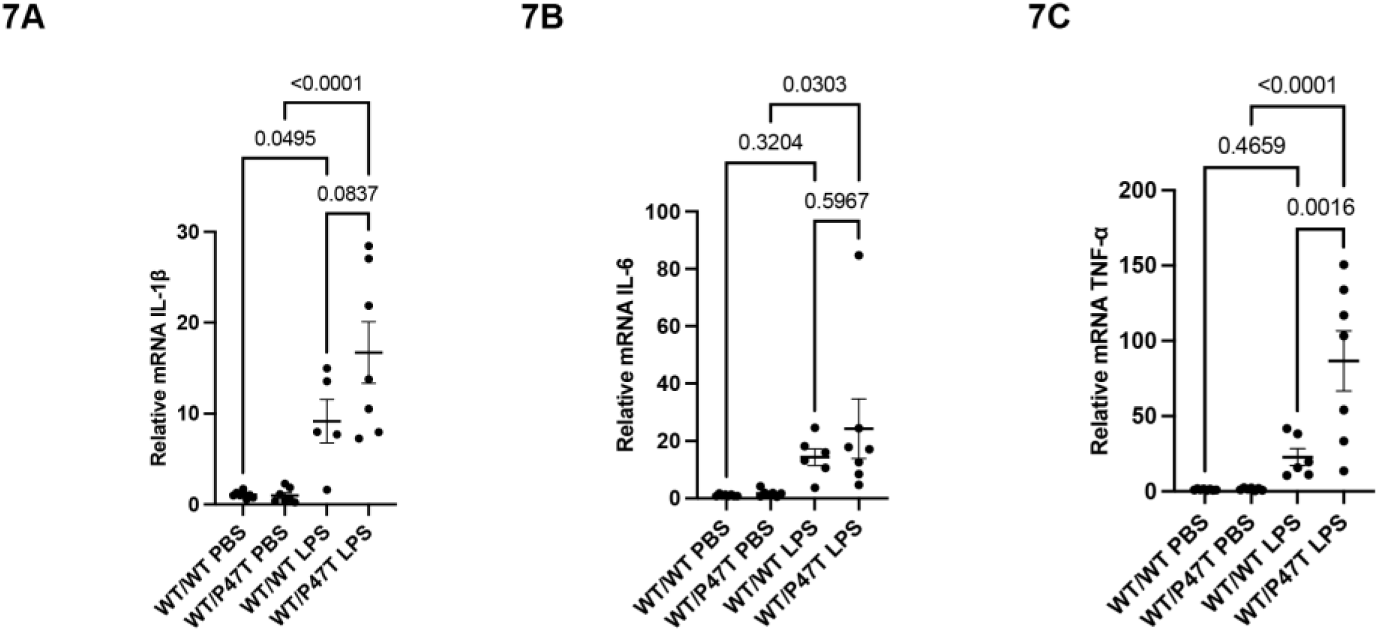
Quantitative PCR is utilized to measure cortical pro-inflammatorycytokines. Both Wwox genotypes demonstrate **a** significant increase in cortical IL-1β in response to LPS. There is a trend towards a more robust reponse in *Wwox ^WT/P47T^* LPS group compared to Iha *Wwox ^WT/WT^* LPS group (p=0.0837) (**7A**). While there is no difference in IL-6 mRNA between LPS groups, there is a significant LPS response in the *Wwox ^WT/P47T^* LPS vs PBS control group (p=0.0303) **(7B).** There is a robust increase in TNF-α in the *Wwox ^WT/P47T^* LPS group compared to PBS control (p<0.0001) and a significant increase in TNF-a between the Wwox wr"""LPS group and the *Wwox ^WT/P47T^* LPS group (p=0.0016) (**7C**).

### Partial Loss of Functional Wwox May Increase LPS-Induced Apoptosis

Apoptosis and specifically neuronal apoptosis were measured using immunohistochemistry stains of NeuN and CC3. There was no difference in area of CC3 or the percent of neurons positive for CC3 in the cortex in response to LPS (**Figure 8A-C**). In the hippocampus, there was an increase in area of CC3+ immunostaining in the *Wwox ^WT/P47T^* LPS group compared to the *Wwox ^WT/WT^* LPS group (p=0.0226) but no difference in CC3-positive neurons (**Figure 9A-C**). This suggests a regional difference in susceptibility to LPS between genotypes that is not affecting neurons.

**Figure 8.**
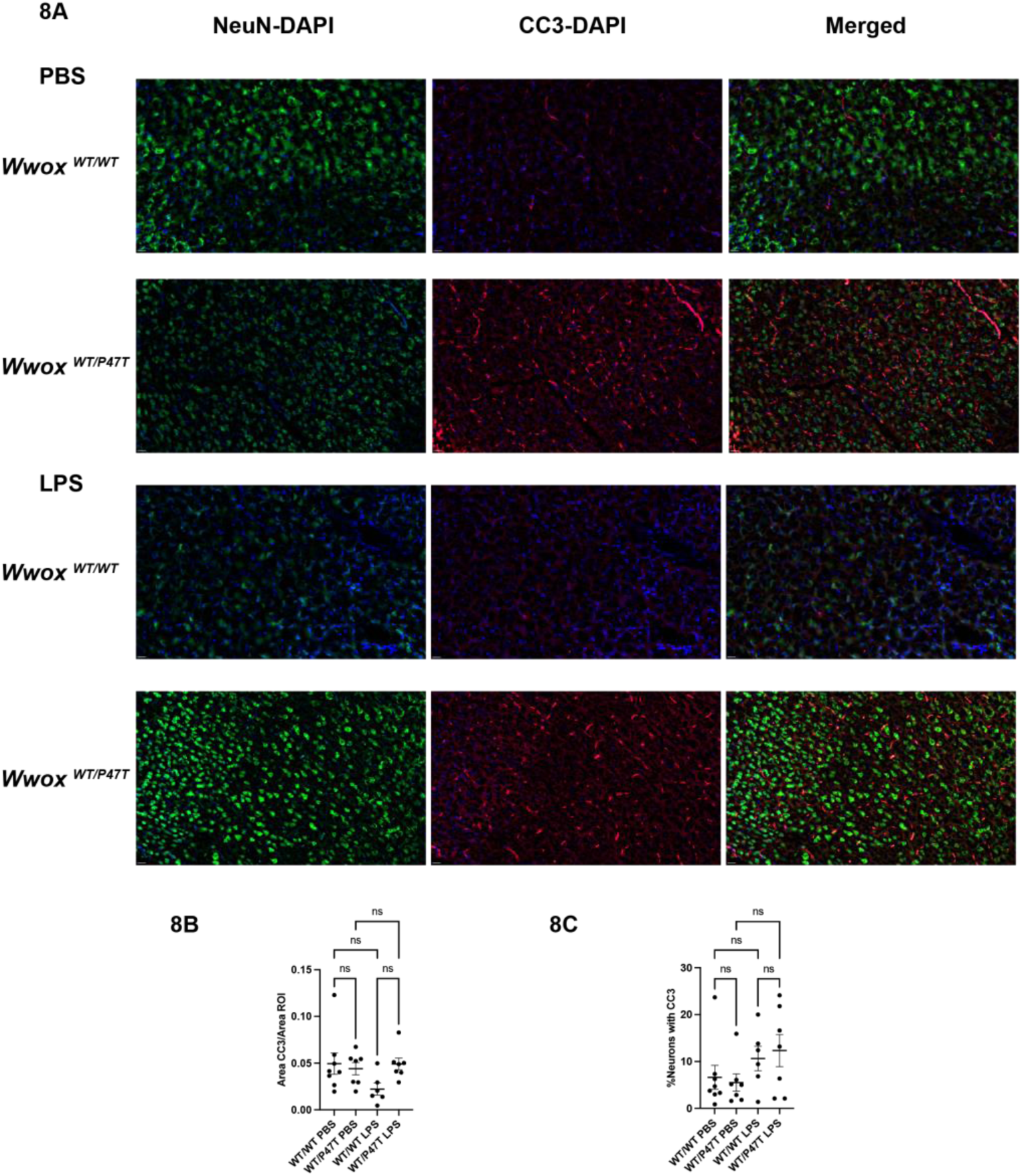
Immunohistochemistry is utilized to analyze presence of apoptosis in the cortex using Cleaved Caspase 3 (CC3) and is costained with Neuronal Nuclei (NeuN). Images are presented with a scale bar at 20μm **(8A).** There is no difference in Area of CC3/Area of ROI ratio **(8B).** There is no difference in percent of neurons positive for CC3 (**8C**).

**Figure 9.**
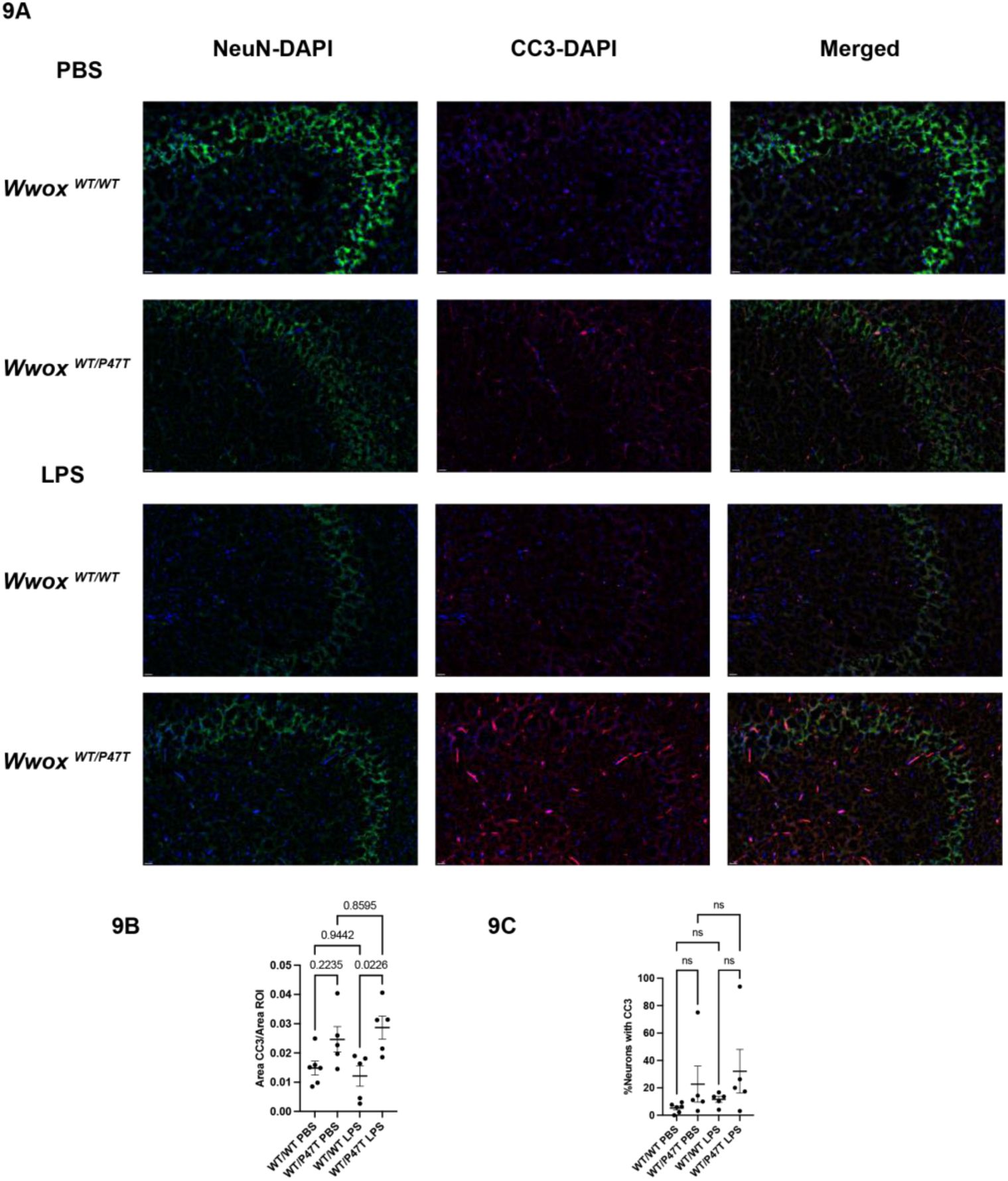
Immunohistochemistryis used to access apoptosis in the hippocampus by using Cleaved Caspase 3 (CC3) and co-stained with Neuronal Nuclei (NeuN). lmagea are presented with scale bar at 20μm **(9A).** There is an increase in area of CC3/area of ROI in the *Wwox ^WT/P47T^* LPS group compared to the *Wwox ^WT/WT^* LPS group (p=0.0226) **(9B).** There is no difference in CC3 positive neurons between groups **(9C).**

### Partial Loss of Functional Wwox Has Variable Impact on LPS-Induced Peripheral Cytokine Expression

Plasma IL-6 levels were significantly elevated in *Wwox ^WT/WT^* LPS mice compared to PBS control (p=0.0168) with no difference in the *Wwox ^WT/P47T^* mice. Plasma TNF-α levels were elevated in the *Wwox ^WT/WT^* LPS mice compared to PBS control (p=0.0025) with no difference in the *Wwox ^WT/P47T^* groups. Both *Wwox ^WT/WT^* (p<0.0001) and *Wwox ^WT/P47T^* mice (p=0.0027) demonstrate an increase in MCP-1. The *Wwox ^WT/P47T^* demonstrate an increase in plasma IFN-y in response to LPS (p=0.0430). There is no between group differences in plasma IL-12p70 or plasma IL-10 (**Supplemental Figure 3A-3F**).

## Discussion

We have investigated the potential role of Wwox in sepsis mortality and encephalopathy by utilizing *Wwox ^WT/WT^* and *Wwox ^WT/P47T^* mice in an LPS sepsis model. Both genotypes demonstrated sickness behavior with reduced distance traveled post-LPS treatment in an open field maze, but the *Wwox ^WT/P47T^* mice also demonstrate reduced time in center. The clinical assessments of mice performed post-LPS treatment demonstrated a significantly accelerated course in sepsis severity in the *Wwox ^WT/P47T^* mice compared to *Wwox ^WT/WT^* mice. *Wwox ^WT/P47T^* mice also demonstrated more severe neuroinflammation in response to LPS. This was evidenced by a statstically signifcant increase in density of microglia and astrocytes in the *Wwox ^WT/P47T^* LPS groups. While both groups demonstrated an LPS effect with an increase in cortical IL-1β mRNA levels, the *Wwox ^WT/P47T^* LPS mice also demonstrated an increase in IL-6 and TNF-α mRNA expression. In terms of apoptosis, the *Wwox ^WT/P47T^* LPS mice showed an increase in brain cleaved caspase 3 immunostaining which was not noted in the *Wwox ^WT/WT^* LPS mice. This apoptosis does not appear to affect neurons and the affected cell type remains to be determined. We note that the difference in apoptosis occurs in the hippocampus rather than the cortex. This may be due to different regions of the brain experiencing suspectibility to LPS at different time points. Lastly, plasma protein expression of inflammatory cytokines differs from mRNA expression of pro-inflammatory cytokines in the brain. This may be due to timing vs difference in technique. Given the possibility of baseline inflammation observed in the *Wwox ^WT/P47T^* mice, future studies may benefit from including a time course and measurement of both mRNA and protein expression of pro-inflammatory cytokines in the brain and plasma.

Overall, this study suggests reduced expression of Wwox is associated with increased sepsis severity and neuroinflammation. GSEA of *Wwox ^P47T/P47T^* brain samples indicated an increase in TNF-α signaling compared to *Wwox ^WT/WT^* littermate controls^13^. Consistent with these findings, our data shows a robust increase in cortical *TNF-α* mRNA in the *Wwox ^WT/P47T^* LPS mice that is not observed in the *Wwox ^WT/WT^* LPS mice. Prior studies suggest, Wwox is essential for TNF-α/ p53 mediated cell death with loss of Wwox increasing hyperactivation of NF-κB causing increased transcription of TNF-α ^21^. Reduced expression of Wwox may mediate sepsis susceptibility via these mechanisms. In addition, loss of endothelial Wwox is associated with increased vascular permeability which may increase the risk of neuroinflammation and contribute to these findings^11,12^.

Overall, our results contribute to prior studies demonstrating *Wwox ^WT/P47T^* mice do not differ significantly from *Wwox ^WT/WT^* littermates at baseline^17^. However, when challenged with experimental sepsis, the *Wwox ^WT/P47T^* mice demonstrate a more severe phenotype, suggesting that loss on one *Wwox* allele is sufficient to increase response to injurious stimuli in the appropriate setting. This is particularly relevant since the expression of WWOX is highly sensitive to environmental stressors, which enhances the general population’s susceptibility to sepsis making this a target for potential intervention^8^. There are several limitations to this study including lack of time course which would better determine the kinetics of inflammatory cytokines in response to LPS. which may better elucidate the timing of peripheral inflammation vs neuroinflammation. The role of blood brain barrier dysfunction was not examined and in future studies we plan to quantify permeability and identify the timing severity of potential LPS-induced blood brain barrier dysfunction. In addition, understating the specific signaling pathways is beyond the scope of this preliminary study and further mechanistic studies will also be conducted.

## Conclusion

In this study we identify that a partial loss of *Wwox* expression increases disease severity and neuroinflammation in experimental sepsis with an increase in sickness behavior, pro-inflammatory cortical cytokines, gliosis, and apoptosis. Given the increased risk of partial loss of Wwox function in response to environmental stressors, this study is applicable to the population at large. Further work is needed to determine the impact of normalizing Wwox expression in setting of sepsis

## Abbreviations

*WWOX*: WW domain-containing oxioreductase
SAE: Sepsis-associated encephalopathy
SCAR 12: Spinocerebellar ataxia type 12
WOREE: Wwox-related epileptic encephalopathy
AD: Alzheimer’s Disease
GWAS: Genome wide association studies
ARDS: Acute respiratory distress syndrome
ip: Intraperitoneal
PBS: Phosphate Buffered Saline
LPS: Lipopolysaccharide
OFT: Open field testing
MSS: Murine sepsis severity score
PBST: Phosphate Buffered Saline-Triton X-100
Iba1: Ionized calcium-binding adaptor molecule 1
GFAP: Glial Fibrillary Acidic Protein
NeuN: Neuron-Specific Nuclear Protein
CC3: Cleaved Caspase 3

## Funding

T32-HL091816 to PD, NIH/NHLBI R01HL127342, R01HL111656, and R01HL158108 to RFM

The authors declare no competing interests

## Author Contributions

PD: Designed and performed experiments, data analysis, manuscript writing, AL-assisted with PCR experiments, MG-assisted with animal colony management, AF-managed animal colonies, TC-managed animal colonies, PS-assisted with flow cytometry experiments, CL-assisted with design of immunohistochemistry experiments, HT-assisted with flow cytometry experiments, manuscript writing, AO-assisted with design of immunohistochemistry experiments, CA-developed transgenic mice, manuscript writing, RM-experimental designing, manuscript writing, and correspondence.

## Acknowledgement

This work was supported by the Behavioral Phenotyping Core (BPC) at the Indiana University School of Medicine

---

**Supplemental Figure 1.**
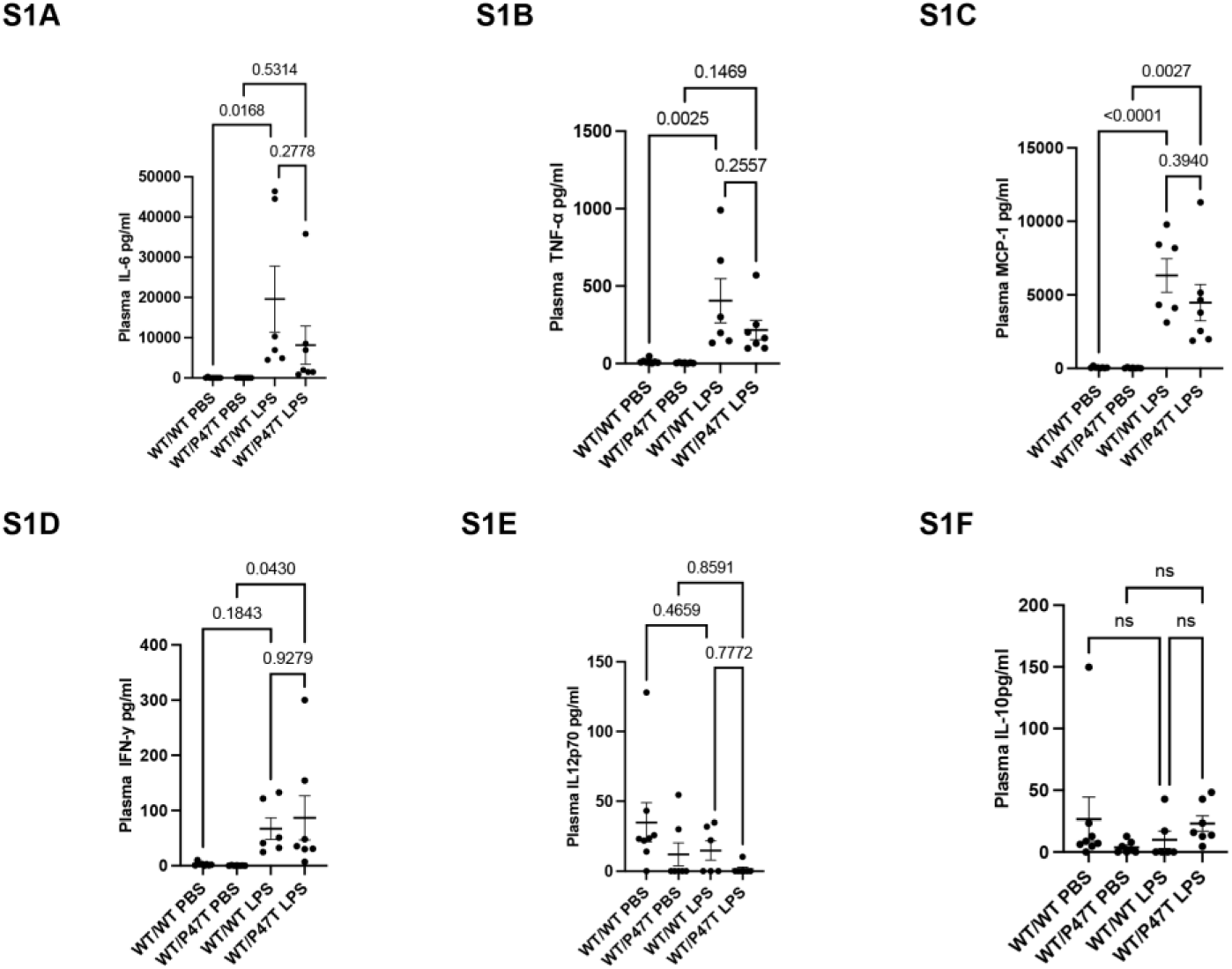
Plasma cytokines were measured using cytometric bead assay.*Wwox ^WT/WT^* LPS group demonstrates a more significant increase in peripheral IL-6 and TNF-α compared to PBS control (p=0.0168) and (p=0.0025), respectively. The *Wwox ^WT/P47T^* did not demonstete a significant difference in II-6 or TNF-α expression in response to LPS **(S1A-81B).** Both *Wwox ^WT/WT^.* (p= <0.0001) and *Wwox ^WT/P47T^* (p=0.0027) demonstrated an increase in plasma IFN-γ expression in response to LPS without a between group difference (S1C). *Wwox ^WT/WT^* demonstrate an increase in plasma IFN-Y in response to LPS (p=0.0430) while there was no significant LPS effect in the *Wwox W1IWT* groups. There was no between group or LPS affect noted in both genotypes with respect to IL-12p70 and IL-10 expression (**S1E-S1F**).

## Notes

### Competing Interest Statement

The authors have declared no competing interest.

